# TrkB-dependent regulation of molecular signaling across septal cell types

**DOI:** 10.1101/2023.06.29.547069

**Authors:** Lionel A. Rodriguez, Matthew Nguyen Tran, Renee Garcia-Flores, Elizabeth A. Pattie, Heena R. Divecha, Sun Hong Kim, Joo Heon Shin, Yong Kyu Lee, Carly Montoya, Andrew E. Jaffe, Leonardo Collado-Torres, Stephanie C. Page, Keri Martinowich

## Abstract

The lateral septum (LS), a GABAergic structure located in the basal forebrain, is implicated in social behavior, learning and memory. We previously demonstrated that expression of tropomyosin kinase receptor B (TrkB) in LS neurons is required for social novelty recognition. To better understand molecular mechanisms by which TrkB signaling controls behavior, we locally knocked down TrkB in LS and used bulk RNA-sequencing to identify changes in gene expression downstream of TrkB. TrkB knockdown induces upregulation of genes associated with inflammation and immune responses, and downregulation of genes associated with synaptic signaling and plasticity. Next, we generated one of the first atlases of molecular profiles for LS cell types using single nucleus RNA-sequencing (snRNA-seq). We identified markers for the septum broadly, and the LS specifically, as well as for all neuronal cell types. We then investigated whether the differentially expressed genes (DEGs) induced by TrkB knockdown map to specific LS cell types. Enrichment testing identified that downregulated DEGs are broadly expressed across neuronal clusters. Enrichment analyses of these DEGs demonstrated that downregulated genes are uniquely expressed in the LS, and associated with either synaptic plasticity or neurodevelopmental disorders. Upregulated genes are enriched in LS microglia, associated with immune response and inflammation, and linked to both neurodegenerative disease and neuropsychiatric disorders. In addition, many of these genes are implicated in regulating social behaviors. In summary, the findings implicate TrkB signaling in the LS as a critical regulator of gene networks associated with psychiatric disorders that display social deficits, including schizophrenia and autism, and with neurodegenerative diseases, including Alzheimer’s.

## INTRODUCTION

The lateral septum (LS) is a GABAergic region in the basal forebrain that integrates sensory and contextual information to influence affective and motivated behaviors ^1,2^. It is implicated in social behaviors ^2–4^, movement and navigation ^5,6^, as well as aspects of both fear ^7,8^ and reward learning ^9,10^. The LS is spatially complex; it spans nearly 2 millimeters across the rostral-caudal axis in the mouse ^5^ and rat brain ^11^, and is divided into dorsal, intermediate, and ventral subregions ^1,12^.

The LS is neurochemically diverse containing predominantly GABAergic neurons that express a variety of inhibitory neuronal markers and neuropeptides including somatostatin, calbindin, calretinin and neurotensin ^12,13^, but sparse labeling of parvalbumin ^14,15^. The LS receives dense innervation from peptidergic hypothalamic systems ^11,16^, and contains spatially defined populations of neurons that express receptors for oxytocin (*Oxtr*), vasopressin (*Avpr1a*) and corticotropin releasing hormone (*Crhr1 and Crhr2*) ^2^. The LS also receives dopaminergic ^11,17^, noradrenergic ^11^, and serotonergic ^18^ innervation. Although recent work has begun to investigate the transcriptional profiles and developmental lineage of neurons in the LS ^19^, most previous studies investigating molecular heterogeneity did not have the methodological capacity to provide comprehensive details on molecular and transcriptional diversity across LS cell types.

LS neurons abundantly express tropomyosin receptor kinase B (TrkB), the receptor for brain-derived neurotrophic factor (BDNF) ^4,20^. While GABAergic neurons generally do not synthesize BDNF, they do highly depend on BDNF, and downstream TrkB signaling in these neurons plays a critical role in supporting GABAergic neuron function ^21–24^. We recently provided evidence that BDNF-producing neurons in the basolateral amygdala project to the LS to act on TrkB-expressing GABAergic neurons in the LS, and that signaling downstream of TrkB in the LS controls social novelty behavior ^4^. While these data implicate TrkB signaling in the LS in controlling social behavior, the molecular mechanisms that mediate this behavior downstream of TrkB remain unknown.

To better understand how TrkB signaling affects LS neurons, we used molecular genetic viral targeting strategies to selectively knockdown TrkB expression in mouse LS neurons. We then used RNA sequencing (RNA-seq) to probe how loss of TrkB in LS neurons alters downstream molecular pathways. To provide cell type-specific context, we mapped genes that were differentially expressed after TrkB knockdown across molecular profiles of LS cell types. To do this, we first generated one of the first comprehensive molecular atlases of gene expression profiles for individual LS cell types in the mouse using single nucleus RNA-sequencing (snRNA-seq). We characterized cell types in the LS, identifying unique markers for the LS broadly and its neuronal subpopulations. We also generated an interactive web application for further exploration of this data to advance understanding of gene expression heterogeneity across LS cell types. Mapping the genes that were regulated after TrkB knockdown across LS cell types identified strong enrichment in microglia and in distinct neuronal subpopulations. Interestingly, upregulated genes were generally enriched in microglia and associated with immune and inflammation pathways, while downregulated genes were broadly expressed across LS neuronal clusters and associated with synaptic signaling and plasticity. Intersecting these datasets also revealed that many of these genes are linked to social behavior and/or BDNF-TrkB signaling. Finally, many of these genes are associated with risk for neurodegenerative disease, such as Alzheimer’s Disease (AD), and with neurodevelopmental disorders that commonly feature deficits in social behaviors, such as schizophrenia (SCZ) and autism spectrum disorders (ASDs).

## MATERIALS AND METHODS

### Animals

Male mice carrying a *loxP*-flanked TrkB allele (strain fB/fB, referenced in text as TrkB^*fl/fl* 25,26^) were bred in our animal facility and used for generating bulk RNA-seq data. TrkB^*fl/fl*^ mice were maintained on a C57BL/6J background. Wild-type male and female mice (C57BL/6J; stock # 000664) were purchased directly from The Jackson Laboratory and used for snRNA-seq experiments. Mice were housed in a temperature-controlled environment with a 12:12 light/dark cycle and *ad libitum* access to food and water. All experimental animal procedures were approved by the Johns Hopkins University School of Medicine Institutional Animal Care and Use Committee.

### Stereotaxic surgeries for delivery of adeno-associated viruses

Adult male mice (10-12 weeks-old) were anesthetized with a mixture of ketamine/xylazine (ketamine, 100 mg/kg; xylazine, 10 mg/kg; intraperitoneal injection). Stereotaxic surgeries were performed using a stereotaxic apparatus (David Kopf Instruments) and Nanoject III injector (Drummond Scientific). A craniotomy was performed and adeno-associated virus (AAV) was injected bilaterally into the LS (+0.90 anterior-posterior (AP), ±0.63 medial-lateral (ML), -3.45 dorsal-ventral (DV)) at 2 nL/s through a pulled glass micropipette. AAV5-EF1a:EGFP was packaged in serotype AAV5 by the University of Pennsylvania Gene Therapy Vector Core; AAV5-EF1a:cre was packaged in serotype AAV5 at the University of North Carolina Vector Core. Titers of AAV viruses were adjusted to 3.0 × 10^12^ particles/mL before injections. After each viral injection, the glass micropipette was left in place for 10 minutes, then slowly withdrawn, and the skin was sutured closed. Mice recovered on a heat pad before returning to their home cage, and animals were used for experiments 4 weeks after surgeries.

### Bulk RNA-seq of LS tissue

Brains were extracted and flash-frozen in 2-methylbutane and then sliced on a mouse brain matrix (Zivic Instruments) to generate two coronal brain slabs (1 mm each) at the level of the LS. From each slab the LS was dissected using a scalpel on a flat metal surface placed over wet ice. Samples were transferred to tubes and placed on dry ice and stored at −80°C until further processing. Total RNA was extracted from samples using Trizol followed by purification with an RNeasy Micro kit (Qiagen). Paired-end strand-specific sequencing libraries were prepared from 40 ng total RNA using the TruSeq Stranded Total RNA Ribo-Zero Gold Library Preparation kit (Illumina). Following quality control on a BioAnalyzer (Agilent), libraries were sequenced on an Illumina HiSeq 3000. The Illumina Real Time Analysis (RTA) module was used for image analysis and base calling, and the BCL converter was used to generate sequence reads, producing a mean of 135 and a range of 35 to 482 million 100-bp paired-end reads per sample.

### Bulk RNA-seq analysis

Gene filtering was carried out based on reads per kilobase per million mapped reads (RPKM) values using the `getRPKM()` function from the *recount* v1.24.1 ^27^ package, where only genes with a mean RPKM greater than 0.1 across all samples were retained. The calculation of the normalization factors was performed using `calcNormFactors()`from *edgeR* v3.40.2 ^28^.

For identification of differentially expressed genes the model matrix was built using the `model.matrix()` while adjusting for total reads assigned to genes and total mapped reads. The differential expression analysis was performed with the *limma* v3.54.2 ^29^ package and `voom()` function. To account for multiple testing, p-values were adjusted by the false discovery rate (FDR) Benjamini-Hochberg algorithm ^30^ using the normalization factors previously calculated. Finally, the differentially expressed genes were extracted based on a FDR < 0.05 threshold.

The differential expression analysis was followed by a gene ontology enrichment analysis with the Gene Ontology: Biological Process (GO:BP), Gene Ontology: Molecular Function (GO:MF) and Kyoto Encyclopedia of Genes and Genomes (KEGG) datasets for each group of differentially expressed genes (upregulated and downregulated). The `compareCluster(pAdjustMethod = “BH”, pvalueCutoff = 0.1, qvalueCutoff = 0.05)` function from the *clusterProfiler* v4.6.2 ^31^ package was used to perform the enrichment analysis, using as the universe dataset the genes that passed the RPKM filter mentioned above. The adjusted p-value was calculated using the Benjamini-Hochberg method for FDR control, and the enriched terms were filtered based on a p-value cutoff of 0.1 and a q-value cutoff of 0.05.

### snRNA-seq of LS tissue

LS dissections were performed as described above. Tissue was pooled from both hemispheres of the LS from 2 mice of the same sex - these 4 pooled LS dissections constituting an individual sample (e.g. N=4 total samples, n= 8 mice; 4 female, 4 male). Nuclei were isolated using the “Frankenstein” nuclei isolation protocol for frozen tissues as previously described ^32,33^. Tissue was homogenized in chilled nuclei EZ lysis buffer (Millipore Sigma #NUC101) using glass dounce with ∼15 strokes per pestle. Homogenate was filtered, centrifuged and resuspended in EZ lysis buffer, followed by three washes in wash/resuspension buffer (1x PBS, 1% BSA, 0.2U/μL RNase Inhibitor (Promega)) and labeled with propidium iodide (PI, ThermoFisher, 1:500). For each sample, 9000 PI+ nuclei were sorted on a Bio-Rad S3e Cell Sorter directly into 23.1μL of reverse transcription reagents from the 10x Genomics Single Cell 3’ Reagents v3.1 kit (without enzyme); water and reverse transcriptase were added to bring each reaction volume to a total of 75μL. The 10x Chromium process was performed to prepare libraries according to the manufacturer protocols (10x Genomics) and sequenced on an Illumina NovaSeq 6000.

### snRNA-seq data analysis

FASTQ data for the n=4 snRNA-seq sample libraries were aligned to the mouse genome (GRCm38/mm10, Ensembl release 98), using 10x Genomics’ software, `cellranger count` (version 6.1), with the `–-include-introns` flag to account for the nuclear transcriptome setting. Raw feature-barcode files were analyzed in R, using the Bioconductor suite of single-cell analytical packages ^34^, as follows: nuclei calling were performed, using `emptyDrops()` on the raw feature-barcode matrices ^35^, using a sample data-driven `lower` threshold. For one particular lower-quality sample (‘2F’), we intersected the thresholded `emptyDrops’` FDR-corrected p-values (FDR < 0.001) with the top barcodes above at computed inflection point (= 6491 total UMIs), as nearly 20,000 barcodes still passed the adaptive approach with a sample-specific `lower` threshold, whereas the known input `n` nuclei = 9,000. This restriction yielded 3,982 nuclei called for sample ‘2F’, compared to 6,728, 6,961, and 7,856 nuclei called from the adaptive `emptyDrops` approach for samples ‘1M’, ‘3M’, and ‘4F’, respectively. We then performed mitochondrial mapping rate adaptive thresholding (using a 3x median absolute deviation from the median, or 3x MAD, to threshold each sample), and filtered out called nuclei with rates higher than the corresponding threshold ^36^. For feature selection, we computed deviance residuals and used total deviance to identify the top 2,000 highly deviant genes (HDGs), followed by PCA in this feature space (i.e. generalized linear models-based PCA, or GLM-PCA)^37^. On the resulting top 50 PCs, we performed graph-based clustering, using `k=20` nearest neighbors, and the walktrap community detection algorithm ^38^ on the resulting graph, preliminarily yielding 35 clusters. We annotated the clusters based on expression of the mouse orthologs of broad cell class markers from a previous study ^32^.

For cluster marker detection, we employed the approach outlined previously ^32^ to characterize strict pairwise test-significant markers per cluster, in addition to cluster-enriched markers. After surveying the marker results and exploring markers’ spatial distribution in the Allen atlas for mouse ISH data, we manually merged glial/non-neuronal clusters to yield one population per cell type, and additionally sub-clustered the preliminarily LS-specific large neuronal clusters. Finally, we flagged two clusters driven by doublets, based on dual broad cell type marker expression and high distribution of doublet density scores using R package `scDblFinder` ^39^, in addition to one cluster driven by low total `n` transcripts. After dropping these from downstream analyses, this resulted in 33 cell populations. We also used the `iSEE` Bioconductor package to generate an interactive app for data visualization^40^.

‘Pseudo-bulking’ analysis was performed for the nine lateral septum clusters by summing the raw counts for each gene within each cell type cluster across replicates, using `registration_pseudobulk()` from *spatialLIBD* v1.11.9 ^41^ with the default values. PCA was then performed with the default data using `runPCA()` and `plotPCA()` from *scater* v1.26.1 ^42^.

### Gene Set Enrichment Analysis

For the gene set enrichment analysis, the pairwise test (1vs1) and enrichment test (1vsAll) results were reshaped to match the format of the `modeling_results` object that is used as input in *spatialLIBD* v1.11.9 ^41^ functions for gene set enrichment analysis. As a filter, the median of the log-transformed counts was calculated for each cell type cluster, and only genes with a median > 0 were used. For the bulk RNA-seq data, only DEGs with FDR < 0.01 were used. The enrichment was made with the function `gene_set_enrichment()` from *spatialLIBD*, with an FDR cut of 0.05, which is applied to the snRNA-seq data. To plot the results, the function `gene_set_enrichment_plot_complex()` from *spatialDLPFC*^*43*^ was used. For later analyses, a list of marker genes enriched in both differentially expressed groups (upregulated and downregulated) for every cell type cluster was extracted. Then, unique (1) microglia genes in the upregulated set when compared to all broad cell type clusters were identified, as well as (2) downregulated lateral septum markers against neuronal broad cell type clusters and (3) LS specific clusters’ underexpressed genes compared to other LS clusters.

## RESULTS

### Knockdown of TrkB in the mouse LS regulates local gene expression

We followed up on previous findings that local knockdown of TrkB expression in the LS impairs social behavior ^4^ to further investigate how molecular signaling downstream of TrkB impacts LS function. We used the same genetic manipulation strategy as in Rodriguez et al., 2022 to locally knockdown TrkB in the LS ^4^, and then subjected TrkB-depleted LS tissue to bulk RNA-sequencing (RNA-seq). Briefly, for experimental animals, we bilaterally injected a cre recombinase-expressing virus (AAV5-EF1a:cre) into the LS of mice carrying a floxed TrkB allele (TrkB^*fl/fl*^ mice), and for control animals we injected a non-cre-expressing virus (AAV5-EF1a:EYFP) into the LS of TrkB^*fl/fl*^ mice (**Fig. 1A**). This manipulation resulted in knockdown of full-length TrkB expression selectively in the LS in the experimental group, while TrkB expression remained intact in the control group (**Fig. 1A**). Four weeks following viral injections, we micro-dissected LS tissue from experimental (n=4) and control mice (n=4). Total RNA was then isolated and processed for bulk RNA-seq. RNA-seq of LS tissue identified 3,750 differentially expressed genes (DEGs) in the `TrkB knockdown group compared to the control group at an FDR-corrected p-value ≤ 0.05: 3,274 genes` (**Table S1**) were upregulated and 476 genes were downregulated (**Fig. 1B, Table S2**). Gene ontology (GO) enrichment analysis showed that genes upregulated following TrkB knockdown in the LS were associated with immune response mechanisms, including immune receptor activity and cytokine-to-cytokine receptor signaling **(Fig. 1C-D**). Genes that were down regulated following TrkB knockdown in the LS were associated with synaptic signaling and plasticity (**Fig. 1C-D**). KEGG pathway enrichment analysis corroborated the GO analysis findings by demonstrating that downregulated genes are involved in processes such as signaling at dopaminergic synapses, while upregulated genes are involved in processes such as cytokine-cytokine receptor interaction (**Fig. 1E**).

**Figure 1:**
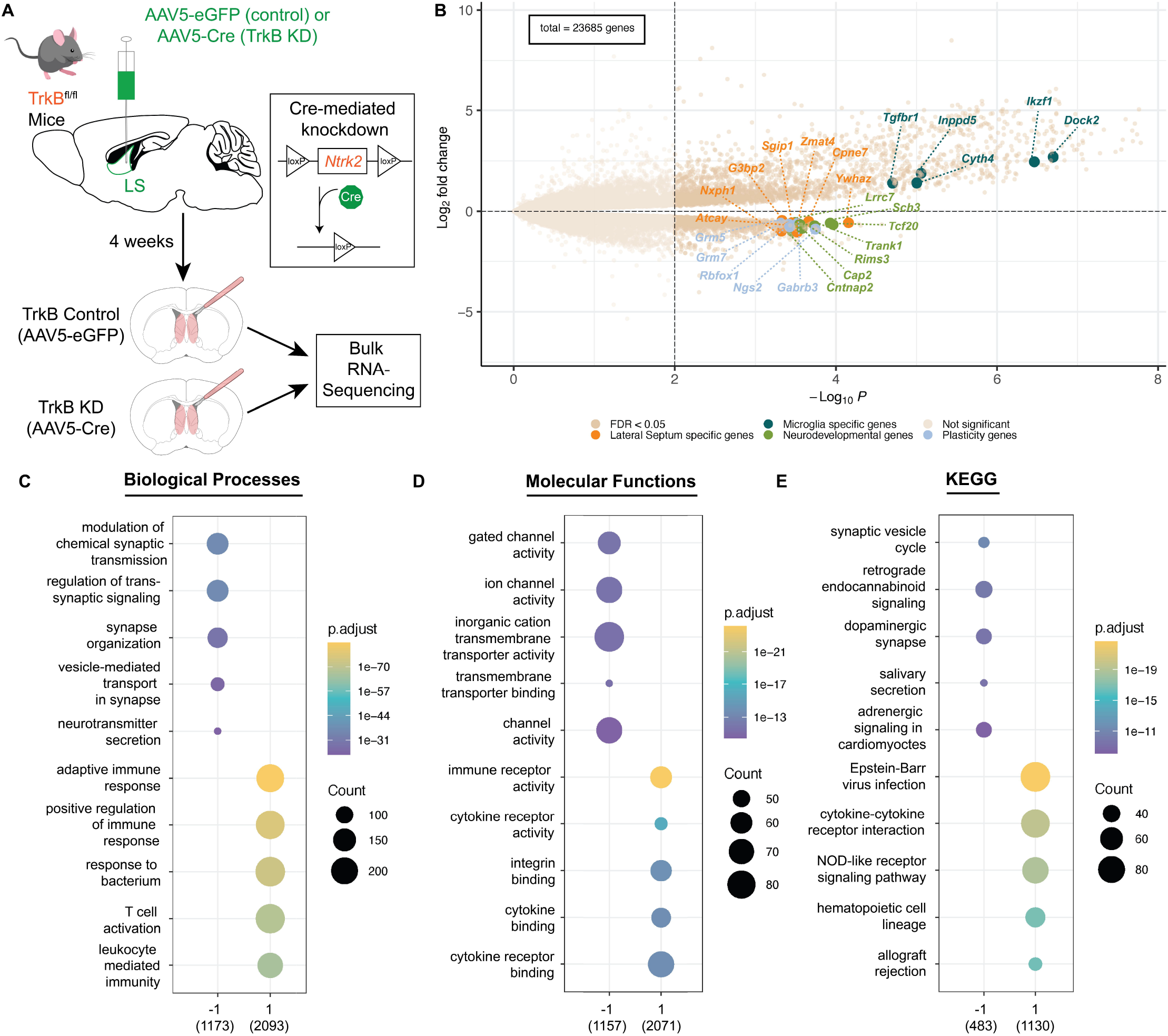
Impact of TrkB knockdown in mouse LS neurons on local gene expression. (**A**) Schematic of viral strategy using cre-mediated recombination to locally knockdown TrkB expression in the LS. (**B**) Volcano plot of differentially expressed genes in bulk RNA-seq of LS tissue samples comparing control versus local TrkB knockdown. Upregulated and downregulated genes were highlighted based on later results (**Fig. 5**). Summaries of gene ontology analysis for (**C**) biological processes, (**D**) molecular functions, and (**E**) Kyoto Encyclopedia of Genes and Genomes (KEGG) annotated pathways.

### Identification of transcriptionally-defined cell types in the mouse LS

To add cell type-specific context to the bulk RNA-seq findings, we generated a molecular atlas of LS cell types using single nucleus RNA-sequencing (snRNA-seq), and then used this data to map TrkB knockdown-induced DEGs across LS cell types. We used the 10x Genomics 3’ single cell platform to perform snRNA-seq on microdissected LS tissue from naive, adult mice (total of N=4 total sequenced samples, each combining the samples of 2 mice of the same sex) (**Fig. 2A**), which yielded 33 data-driven cell types (**Fig. 2B**). To characterize neuronal and non-neuronal populations, we used previously defined broad cell type definitions ^32^ including a pan-neuronal marker (*Snap25*, **Fig. 2C**), markers for GABAergic neurons (*Gad2*, **Fig. 2D**), astrocytes (*Slc1a2*, **Fig. 2E**), oligodendrocytes (*Mbp*, **Fig. 2F**), and choroid plexus ^44,45^, ependymal cells ^46,47^, and differentiating neuroblasts ^48^ (**Fig. 2G**).

**Figure 2:**
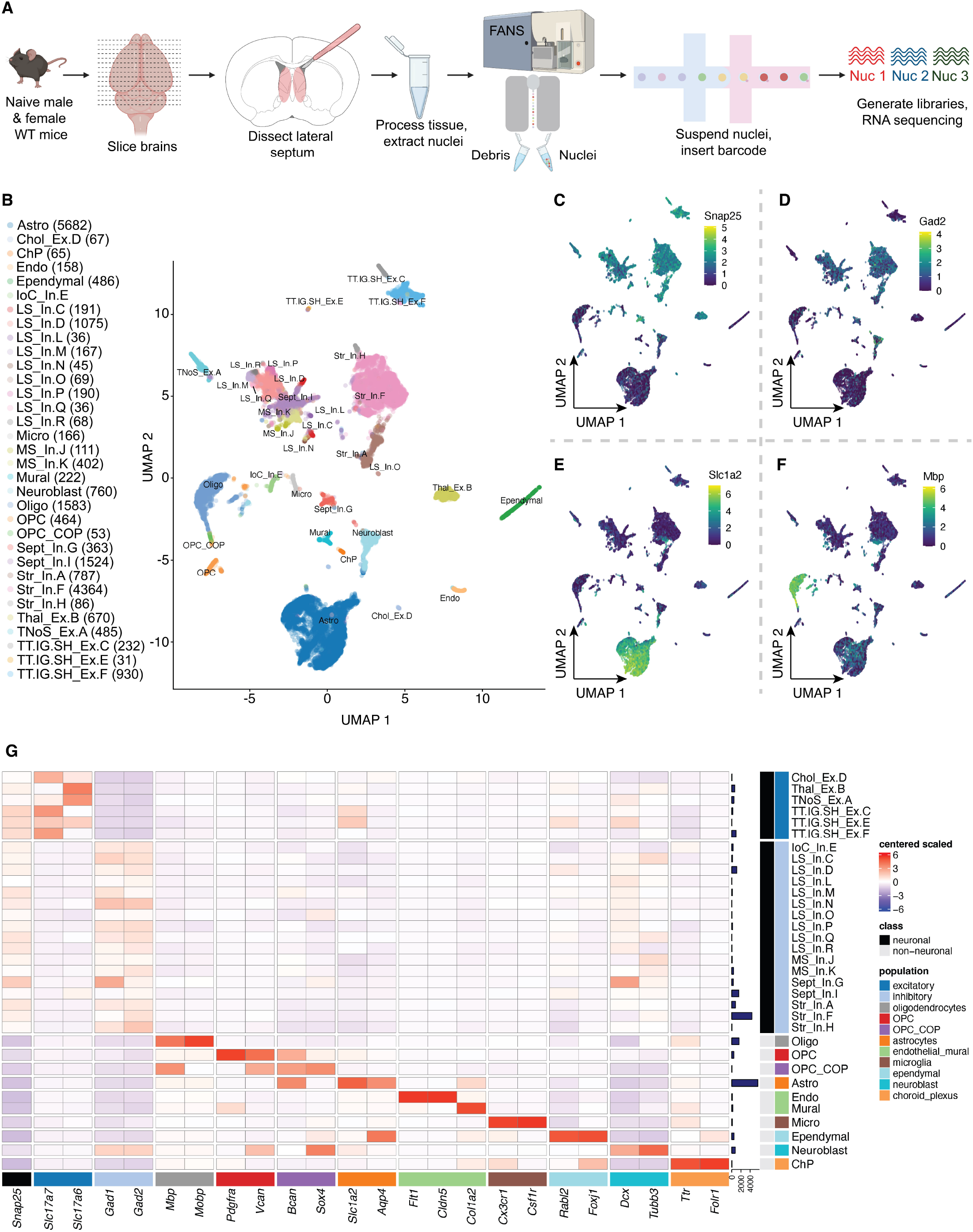
Identification of transcriptionally-defined cell types in mouse LS. (**A**) Schematic of experimental design for snRNA-seq of mouse LS tissue. Tissues from 2 mice of the same sex were pooled together for each individual sample for a total of N=4 samples (generated from n=4 male,4 female mice). (**B**) Uniform manifold approximation and projection (UMAP) of identified cell types, with nuclei counts per clusters. Feature plots for expression of (**C**) *Snap25*, (**D**) *Gad2*, (**E**) *Slc1a2*, and (**F**) *Mbp* across the dataset. (**G**) Heatmap of cell type markers used to characterize the dataset with normalized expression values (logcounts) centered and scaled.

Of the 33 total clusters, 23 were identified as neuronal. To map the anatomical origin of these clusters, we used two Allen Brain Atlas data resources ^49^. First, we used the differential gene expression search function to identify genes with the highest expression in the septum proper and in regions surrounding the septum. We then examined expression of these genes across the neuronal clusters. Second, we identified DEGs that were enriched in each neuronal cluster, using pairwise comparisons, referencing Allen Brain Atlas’ gene expression atlas to characterize the spatial distribution of these DEGs in the regions within and surrounding the septum. The best markers identified for both regions and subregions are summarized as a heatmap of marker expression across all neuronal clusters (**Fig. 3A**). Specifically, these analyses identified *Trpc4* (**Fig. 3B**), *Homer2* (**Fig. 3D**), and *Ptpn3* (**Fig. 3E**) as reliable markers for LS neurons, while *Dgkg* (F**ig. 3C**) is expressed in the septum more broadly.

**Figure 3:**
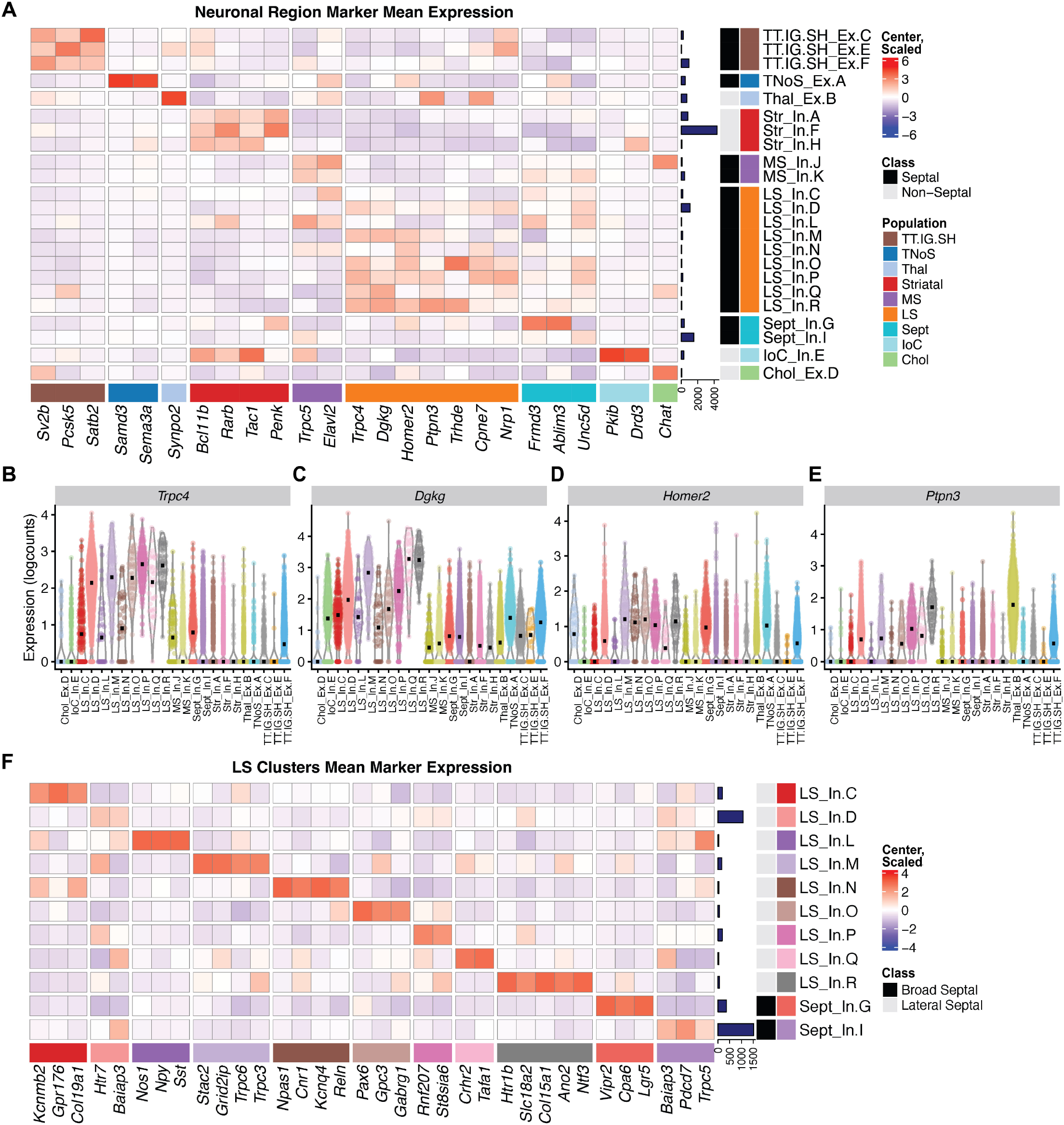
Characterization of neuronal clusters within mouse LS. (**A**) Heatmap of marker genes for various brain regions used to characterize neuronal clusters. Violin plots showing (**B**) *Trpc4*, (**C**) *Dgkg*, (**D**) *Homer2*, and (**E**) *Ptpn3* expression, which were identified as specific markers for LS neurons. (**F**) Heatmap of identified cell type-specific markers for each of the LS neuronal clusters. Heatmaps show normalized expression values (logcounts) centered and scaled.

Of the 23 neuronal clusters, 18 were identified as mapping to the septum, including 9 that mapped to the LS and 3 that mapped to the medial septum (MS). We identified 1 cholinergic population specific to the MS, and 2 that mapped to both the MS and LS, which we labeled as “Septal” clusters. In addition, we identified 3 clusters from the rostrodorsal part of the septum, where the tenia tecta, indusium griseum, and septohippocampal nucleus localize, which cluster together as one population (TT.IG.SH). We also identified 1 cluster from the caudal-medial septal region known as the triangular nucleus of the septum (TNoS). In addition to septal populations, we identified 3 striatal clusters, 1 cluster from the islands of calleja (IoC), and 1 from the thalamus (Thal). We then plotted gene expression of receptors enriched in the septum (**Fig. S1**).

We next molecularly characterized the LS clusters by identifying cell type markers for each LS and septal cluster. Here, we used pairwise *t*-test comparisons across all of the neuronal clusters to identify markers unique to, or most expressed within, each cluster. For LS inhibitory cluster D (LS_In.D) and septal inhibitory cluster I (Sept_In.I), this conservative approach failed to identify any statistically significant unique markers. Thus, to identify genes enriched in clusters LS_In.D and Sept_In.I, we adopted a 1 versus all approach and compared the cluster of interest to the combined gene sets for the rest of the neuronal clusters. A summary of cluster-specific markers is displayed as a heatmap (**Fig. 3F**). We then validated the topographical organization of our clusters (**Fig. S2**) ^50^. Additionally, we performed “pseudo-bulking” by summing the UMI counts from each gene within each individual LS cluster for each sample to generate sample-specific, cluster-enriched expression profiles. Unsupervised principal component analysis of these “pseudo-bulked” data per cluster and sample revealed that the top two principal components (PC) are able to categorize the LS clusters into 3 separate groups (**Fig. 4A**): group one being composed of LS_In.C, LS_In.L, and LS_In.N, group two being composed of LS_In.D, LS_In.M, LS_In.P, LS_In.Q and LS_In.R, and group three being a single cluster (LS_In.O). PC1 compared to PC3 and 4 maintain this separation between group one and two, while PC2 compared to PC3 and 4 maintain this separation of LS_In.O from other clusters, suggesting these clusters may be functionally or spatially distinct. Pairwise and 1vAll markers were computed against our broad clusters. These markers show enrichment in the LS clusters from the respective groups compared to neuronal clusters (**Fig. 4B**) and LS clusters (**Fig. 4C**-**J**).

**Figure 4:**
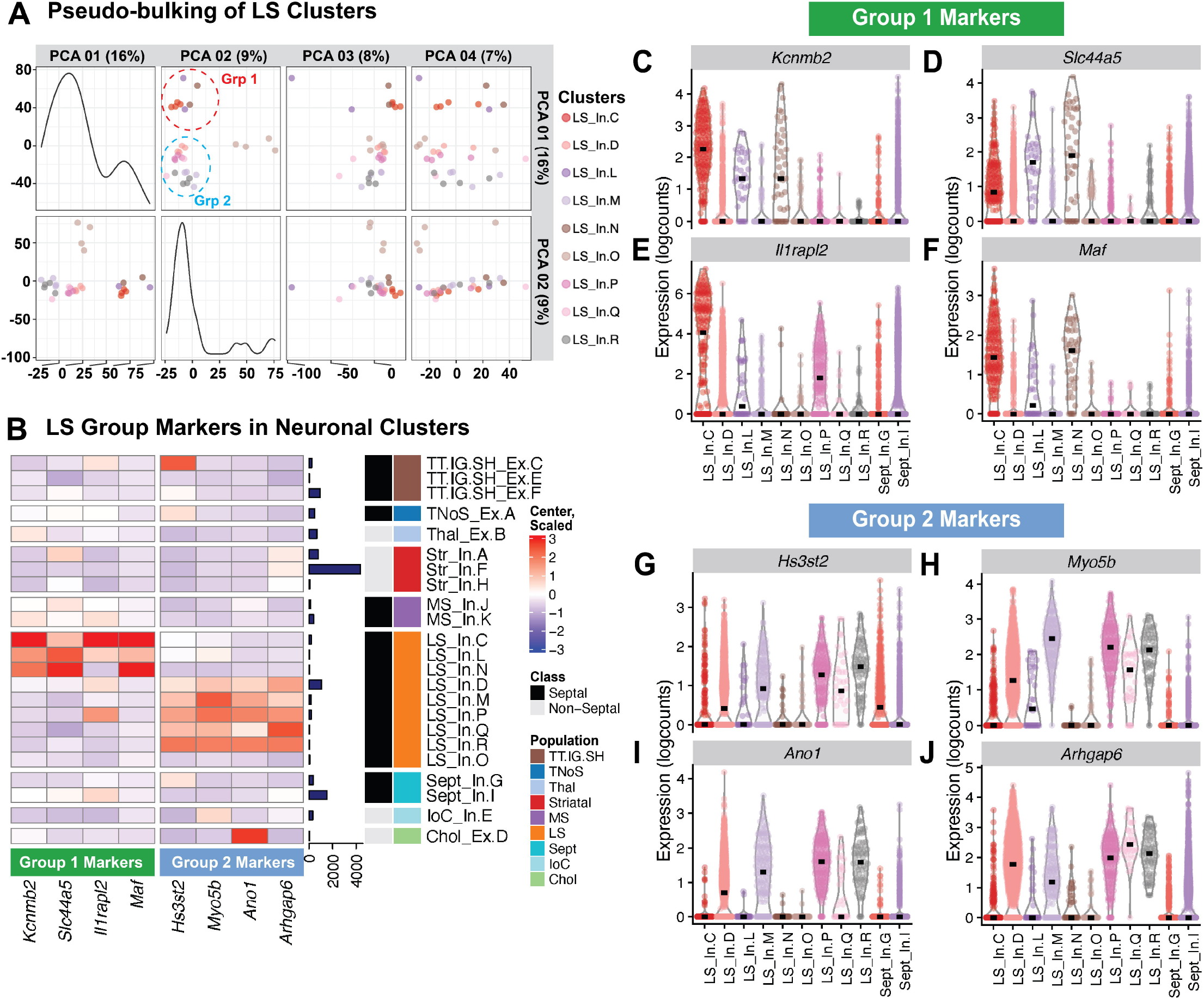
Pseudo-bulking the LS cluster data reveals three groups of neurons. (**A**) Principal Component Analysis of LS clusters pseudo-bulked expression profiles across all samples. (**B**) Heatmap of genes enriched in group 1 (LS_In.C, LS_In.L, and LS_In.N), group 2 (LS_In.D, LS_In.M, LS_In.P, LS_In. Q, and LS_In.R) and previously calculated markers for LS_In.O, Sept_In.G, and Sept_In.I across our neuronal clusters. Normalized expression values (logcounts) centered and scaled. Violin plots of these genes expressed across LS populations for group 1 (**C**-**F**) and group 2 (**G**-**J**) highlight the specificity of these markers.

### Enrichment of genes associated with local TrkB knockdown across LS cell types

To better understand how TrkB knockdown impacts the cellular landscape of the LS, we used a previously validated strategy ^41^,^43^ to map enrichment of DEGs from the TrkB knockdown dataset (FDR < 0.01) across the broad clusters identified in the LS snRNA-seq data (Fisher’s exact test, FDR < 0.05) (**Fig. 5A**). Similarly, we ran an enrichment test using DEGs from the TrkB knockdown data set (FDR < 0.01) across the septal neuronal clusters from the LS snRNA-seq dataset (Fisher’s exact test, FDR < 0.05) (**Fig. 5B**). We then applied an enrichment analysis to identify upregulated genes that are enriched in the microglia broad cluster and downregulated genes enriched in the LS broad cluster compared to other septal clusters, in addition to genes in the LS_In.D and LS_In.O clusters that more significantly express downregulated DEGs from the TrkB knockdown dataset. The enrichment analyses yielded four sets of genes: genes with unique LS expression, genes with unique microglial expression, plasticity genes, and neurodevelopmental genes (**Table S3**). To understand the expression patterns of these four gene sets across the snRNA-seq data, we generated a heatmap showing these gene set expressions across the snRNA-seq broad clusters (**Fig. 5C**) and individual neuronal clusters (**Fig. 5D**). Expression levels of DEGs that are both downregulated following TrkB knockdown and enriched in the LS broad cluster with unique LS expression are highlighted as violin plots (**Fig. 5E**-**L**).

**Figure 5:**
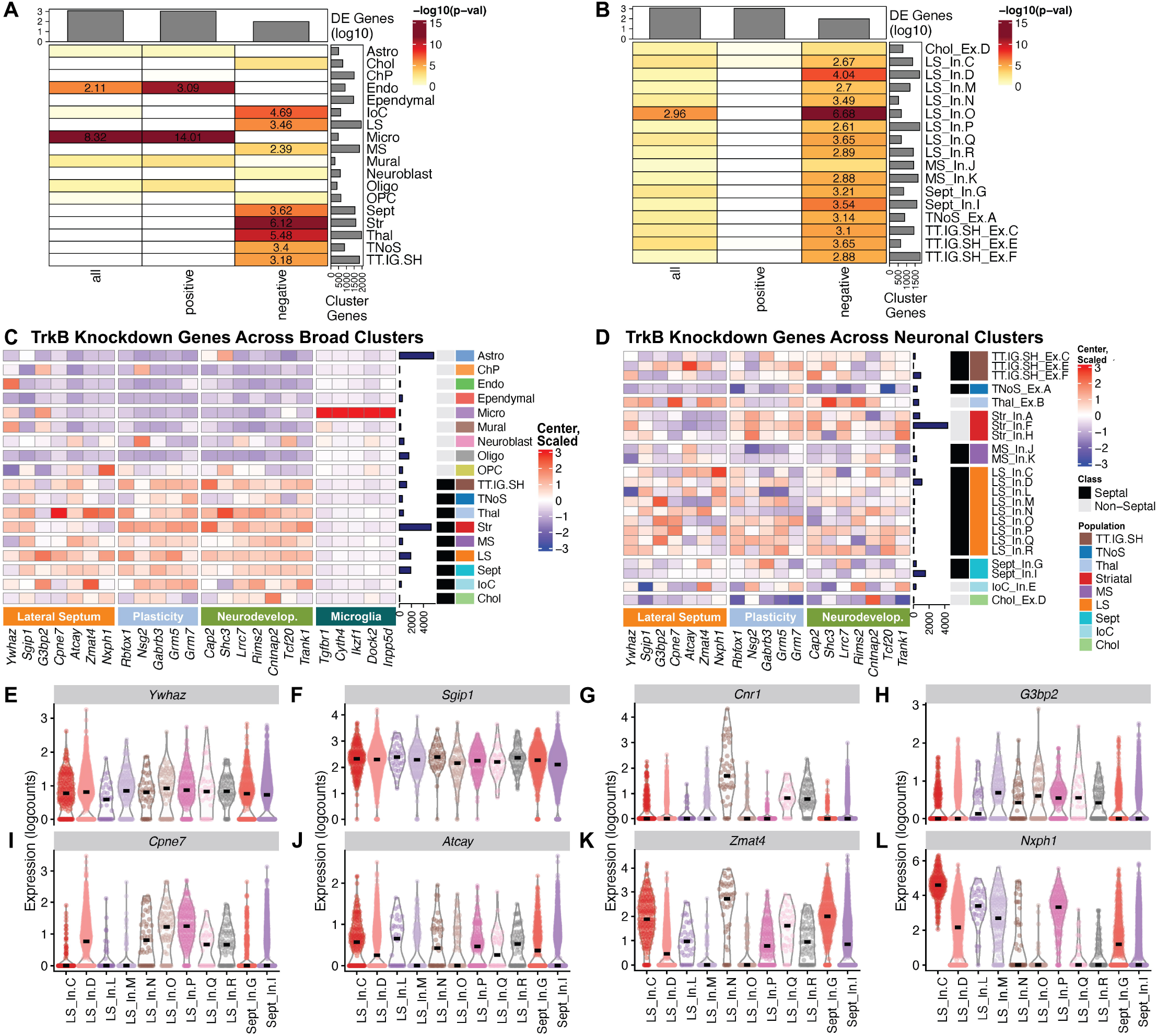
Enrichment of differential expression signals from local TrkB knockdown across snRNA-seq LS dataset. (**A**) Enrichment analysis of all, positive, and negative DEGs from the TrkB knockdown dataset across broad cellular clusters. (**B**) Enrichment analysis of all, positive, and negative DE genes from the TrkB knockdown dataset across all neuronal clusters from the septum. (**C**) Heatmap of DE genes from the TrkB knockdown dataset enriched in lateral septal and microglial broad clusters across broad clusters. DE genes from the TrkB knockdown dataset are divided into genes unique to the LS, plasticity genes, and neurodevelopmental genes. (**D**) Heatmap of DE genes from the TrkB knockdown dataset that are enriched in the lateral septal broad cluster across the neuronal clusters. Heatmaps show normalized expression values (logcounts) centered and scaled. Violin plots of (**E**) *Ywhaz*, (**F**) *Sgip1*, (**G**) *Cnr1*, (**H**) *G3bp2*, (**I**) *Cpne7*, (**J**) *Atcay*, (**K**) *Zmat4*, and (**L**) *Nxph1* expression, genes with unique LS expression identified from downregulated DE genes from the TrkB knockdown dataset enriched in the LS broad cluster, in addition to *Cnr1* expression.

## DISCUSSION

### TrkB knockdown in the LS induces robust changes in synaptic signaling and neuroimmune gene networks

TrkB knockdown resulted in robust depletion of genes associated with synaptic organization and signaling, including vesicular machinery for neurotransmitter release, mechanisms for chemical synapse transmission, as well as various synaptic channels and transporters. This was particularly true for genes associated with dopaminergic synapses, which is notable given that dopamine signaling in the LS controls social behavior ^17,51^. In addition, TrkB knockdown depletes expression of genes associated with retrograde endocannabinoid signaling, which is linked to social behavior ^52^. Conversely, TrkB knockdown increased expression of neuroimmune and neuroinflammatory gene networks, implicating cytokine-cytokine receptor signaling, immune receptor activity, and NOD-like receptor signaling. The interaction between the immune system and brain circuits that control social behavior are complex, but critical for understanding both normal and aberrant social behavior. In macaques, social hierarchy dictates basal level of immune responses and regulates the immune response to lipopolysaccharide (LPS)^53^, which induces cytokine and chemokine expression to trigger immune system responses ^54^. During LPS challenge, higher social status macaques showed an increased antiviral response, while lower social status macaques increased inflammatory gene networks for bacterial infections ^53^. Human subjects experiencing chronic social isolation also experienced acute increases in proinflammatory gene networks and decreases in type-1 interferon signaling ^55^. While the specific mechanisms are not understood, experience can shape immune system function while the immune system can shape behavior ^54^, and our findings suggest that signaling downstream of TrkB in the LS can regulate the interaction between immune responses and synaptic signaling to influence social behavior.

Supporting this idea, immune responses can regulate BDNF-TrkB signaling, which may then influence behavior. Interleukin-1 (IL-1) is a proinflammatory cytokine that is the endogenous agonist of the IL-1 receptor (IL-1R), and in rats, peripheral IL-1R receptor agonists decrease motivation for social interaction, which can be reversed by an IL-1R antagonist ^56^. Scavenging extracellular BDNF from the media of cultured neurons increases proinflammatory cytokines, like TNFα ^57^. Bacterial infection affects memory consolidation in older rats, reduces levels of BDNF in the hippocampus and decreases activated TrkB in the hippocampus, which is rescued by IL-1R antagonists ^58^. In the dorsal root ganglia (DRG), induction of inflammation upregulates TNFα, which in turn upregulates BDNF to impact DRG excitability and hyperalgesia ^59^. Based on these relationships, we hypothesize that regulation of inflammatory/immune networks by TrkB in the LS may contribute to social behavior regulation.

### Identification of unique neuronal cell types in the LS

Comprehensive molecular profiles for all LS cell types are lacking. Our snRNA-seq analysis of the LS revealed 9 different LS cell types, with 2 broad septal clusters that comprise a mix of LS nuclei and nuclei from other septal subregions (**Fig. 3F**). As expected, all LS clusters are GABAergic (**Fig. 2G**)^12,13^. The LS expresses various receptors for neuropeptides like oxytocin and vasopressin ^2^, and monoamine receptors ^49^. We mapped the expression of monoamine receptors (**Fig. S1A**) and various peptides and peptide receptors (**Fig. S1B**) to the neuronal clusters. An interactive web browser of this LS snRNA-seq dataset was also generated for further exploration available at https://research.libd.org/septum_lateral/#interactive-websites ^40^.

Prior to these analyses, no unifying marker for the septum broadly or the LS specifically was established. Our data suggest that *Trpc4, Homer2*, and, to a lesser degree, *Ptpn3* are good LS markers, while *Dgkg* broadly marks the septum. This data is consistent with *in situ* hybridization data from the Allen Brain Atlas ^49^. Unsupervised clustering of “pseudo-bulked” data per cluster and sample reveals 3 different types of LS cell type clusters. LS_In.C, LS_In.L, and LS_In.N group together and LS_In.D, LS_In.M, LS_In.P, LS_In.Q and LS_In.R group together, with LS_In.O being distinct (**Fig. 4A**). Interestingly, the LS marker *Ptpn3* is preferentially expressed in the second group of LS cells (**Fig. 3E**), which prompted us to generate markers selective for these groups (**Fig. 4B**-**J**). Besnard and Leroy recently proposed a topographical map of LS subregions including potential molecular markers (**Fig. S2A**)^50^. We generated a heatmap of these markers across the neuronal clusters to predict their spatial location (**Fig. S2B**). Some LS clusters express markers from a single topographical region, like LS_In.C which expresses markers found in the caudal part of the dorsal LS, whereas clusters like LS_In.L express multiple topographical markers. Notably, *Pax6* is a cell type marker for the LS_In.O cluster and a marker for the ventral LS broadly and LS_In.O also expresses other markers found in the ventral LS.

Neurotensin (Nts)-expressing neurons in the LS control both social and feeding behaviors ^13,60^, but we did not observe robust *Nts* expression in the snRNA-seq data. While the reasons for this are not entirely clear, it may result from technical issues associated with snRNA-seq as similar discrepancies for other genes (e.g. *Pvalb*) have been noted. This may be due to specific transcripts being rapidly shuttled out of the nuclear compartment ^32^. This notion is supported by *Nts* being robustly expressed in the bulk RNA-seq data, where it was downregulated following TrkB knockdown (**Table S2**). Future experiments using other methodologies such as single-molecule fluorescence *in situ* hybridization could delineate in which clusters *Nts* is expressed. *Drd3* expression in the LS is critical for social behavior ^51^, and while we don’t observe *Drd3* expression in LS clusters, *Drd3*-positive neurons in the LS may be transcriptionally similar to *Drd3*-positive neurons of the islands of calleja. The island of calleja cluster robustly expresses the broad septal marker *Dgkg* (**Fig. 3C**), which is not expressed in the islands of calleja according to Allen Brain Atlas ^49^. Similarly, the striatal clusters express *Dgkg;* however we propose that these may be LS neurons or LS-nucleus accumbens boundary neurons, and may explain why the striatal cluster Str_In.F is so large and heterogeneous.

### TrkB knockdown in the LS regulates genes associated with social behavior and neurodevelopmental disorders

To understand how TrkB knockdown impacts unique cell types in the LS, we conducted an enrichment test of DEGs induced by TrkB knockdown across the broad LS clusters (**Fig. 5A**) and within the neuronal septal clusters (**Fig. 5B**). Generally, downregulated genes are associated with synaptic signaling and plasticity, which are genes that are shared by the other broad neuronal (**Fig. 5A**) and individual neuronal (**Fig. 5B**) clusters in the enrichment testing. We conducted multiple set operations to determine whether any downregulated DEGs were enriched in the LS broad cluster and the LS_In.D and LS_In.O clusters. For downregulated genes, LS_In.O was not enriched for any TrkB knockdown downregulated genes when compared to other LS clusters, while LS_In.D contained a single gene - *Ywhaz*, a gene that was also enriched in the LS broad cluster when compared to other septal clusters. *Ywhaz* encodes the protein 14-3-3ζ, and genetic variants of *Ywhaz* are associated with autism spectrum disorders (ASDs) ^61,62^ and schizophrenia ^62,63^. *Ywhaz* knockdown in zebrafish causes social recognition deficits ^64^ similar to those observed in mice after TrkB knockdown in the LS ^4^.

The enrichment analysis of downregulated TrkB genes expressed in the LS broad cluster yielded 42 genes. We examined expression of these genes across all clusters, and researched their role in behavior, characterizing them into 3 groups, the first of which are genes with unique expression in the LS broad cluster. The other gene sets that were enriched in the LS were characterized as genes associated with synaptic plasticity or genes associated with neurodevelopmental and psychiatric disorders (**Table S3**). For genes with unique expression in the LS, the most enriched was *Ywhaz*, which as noted above, regulates social behavior ^64^. The next was *Sgip1*, a gene involved in clathrin mediated endocytosis. SGIP1 interacts with the cannabinoid CB_1_ receptor to alter its signaling ^65^. *Sgip1* knockdown mice show decreased anxiety, enhanced fear extinction, and reduced nociception. *Sgip1* knockdown also enhances withdrawal from THC and causes enhanced binding of THC, a CB_1_ receptor agonist and morphine ^65^. While *Sgip1* is broadly expressed in the LS (**Fig. 5D**), *Cnr1* is only expressed in LS_In.Q, and LS_In.R, and it is a marker for the LS_In.N cluster (**Fig. 3F**). The remaining genes (*G3bp2, Cpne7, Atcay, Zmat4*, and *Nxph1*) of the unique LS genes are generally understudied in their role in regulating behavior, although *Zmat4* is associated with schizophrenia ^66^. The five genes associated with synaptic plasticity are *Rbfox1, Nsg2, Gabrb3, Grm5*, and *Grm7. Rbfox1* is an RNA binding protein that regulates alternative splicing of neuronal transcripts ^67^ including TrkB isoforms to control BDNF-dependent LTP ^68^. *Nsg2* is an AMPA receptor interacting protein ^69^. Both *Rbfox1* and *Nsg2* regulate neuronal excitability and neurotransmission. *Gabrb3* encodes a subunit of the GABA_A_ receptor ^70^, and *Grm5* and *Grm7* encode metabotropic glutamate receptors (mGluR5 and mGluR7). Mouse models with genetic knockdown of these synaptic receptors display social behavior deficits ^70–72^. We identified 7 genes (*Cap2, Shc3, Lrrc7, Rims2, Cntnap2, Tcf20*, and *Trank1*) associated with neurodevelopmental and psychiatric disorders that feature social impairments. *Cntnap2* was among the first genes with evidence for rare and common variants contributing to ASDs ^73^. *CAP2*, a gene that regulates actin dynamics to shape spine density and dendritic complexity, is decreased in the hippocampus of individuals diagnosed with schizophrenia (SCZ)^74^. *Shc3* encodes the ShcC protein, which transmits neurotrophin signals from TrkB to the Ras/MAPK pathway ^75^ and modulates NMDA receptor function ^76^. Genetic variation at the *SHC3* locus is associated with SCZ ^77^. *Lrrc7* encodes Densin-180, which interacts with synaptic receptors, and *Lrrc7* mutant mice have social and mood-related impairments ^78^. Interestingly, these impairments are alleviated by mGluR5 allosteric modulation, and *Grm5* is one of synaptic plasticity genes identified above (**Fig. 5C**-**D**). *Lrrc7* is a risk gene for both ASDs ^79^ and emotional dysregulation in children ^80^, and iPSCs from schizophrenia patients display *de novo* mutations in *Lrrc7* ^*81*^. *De novo* mutations associated with ASDs are found in transcription factor 20 (*Tcf20*) ^82^ and *Tcf20* knockdown mice display impairments in social novelty recognition and pup communication ^83^. Moreover, *Rbfox1*, one of the plasticity genes identified in the enrichment analyses is also linked to risk for ASDs ^84^, *TRANK1* is a susceptibility gene for bipolar disorder ^85^, and *RIMS1* was identified in a PTSD TWAS ^86^. Together, these findings suggest that TrkB knockdown in the LS decreases expression of genes that control synaptic signaling, which may alter LS function to give rise to behavioral impairments that are relevant for disorders where social communication is affected.

### TrkB knockdown induces immune signaling and inflammation relevant to neurodegenerative, neurodevelopmental and neuropsychiatric disorders

The majority of genes upregulated in response to TrkB knockdown are associated with inflammation or immune processes. Since the broad microglia cluster showed significant enrichment for upregulated genes (**Fig. 5A**), we conducted an enrichment analysis that identified 52 DEGs specifically upregulated in this cluster. Genes that were both selective for and highly expressed in the microglial cluster were *Tgfbr1, Cyth4, Ikzf1, Dock2*, and *Inpp5d*, many of which are associated with Alzheimer’s Disease (AD). Neuroinflammation is a hallmark of AD pathology ^87^, which induces proinflammatory signals and increases proteins that maintain proinflammatory states, ultimately leading to degeneration. In response to inflammation, local astrocytes release TGF-β to promote microglial homeostasis and repress inflammatory responses, and *Tgfbr1* encodes one of the receptors for TGF-β. *Dock2* encodes a protein that regulates microglial innate immunity and neuroinflammatory responses, which is elevated in microglia of human AD subjects ^88^. Microglial and astrocyte activation is initially beneficial as it facilitates amyloid-beta clearance, but prolonged production of proinflammatory cytokines promotes degeneration ^89^. *Inpp5d* is an AD risk gene ^90,91^ inhibits signal transduction initiated by immune cell activation ^92^. Its expression is positively associated with amyloid plaque density in the human brain and is localized selectively in plaque-associated microglia ^92^. *Ikzf1* is a microglial gene that controls the morphology and function of microglia that is necessary for synaptic plasticity in the hippocampus ^93^. Ikzf1 is elevated in the hippocampus of mouse models of neurodegeneration and in human AD samples ^93^. BDNF-TrkB signaling is neuroprotective, but is decreased as AD pathology progresses, contributing to degenerative processes ^94,95^. Importantly, BDNF is protective against amyloid-beta toxicity in cultures of septal neurons ^96^, and given findings that TrkB knockdown in the LS promotes inflammation, our findings may provide insight into the interaction between BDNF-TrkB signaling and inflammation during degeneration in the LS.

Neuroinflammation is also hypothesized to play a role in the etiology of neurodevelopmental and neuropsychiatric disorders, including SCZ ^97^. The human *CYTH4* has a primate-specific GTTT-repeat and longer repeats (7-9 vs <7) were found in healthy controls compared to patients with neurodegenerative disorders ^98^ and bipolar disorder ^99^, while extremely long versions (10-11) were seen in SCZ patients ^100^. In a repeated ketamine exposure model of SCZ, decreased expression of *Cyth4* is found in rat cortex ^101^. Neuroinflammation is also linked to ASD ^102,103^. Maternal immune activation increases the risk for ASD ^104,105^ and mouse models of maternal infection lead to deficits in social behaviors ^106,107^. In a study intersecting maternal immune environments in mice with offspring behavior, influenza vaccination improved abnormal fetal brain lamination induced by lipopolysaccharide (LPS) infection, and also prevented deficits in offspring of mothers infected with LPS ^106^. Whole-genome sequencing of these mice suggests that *Ikzf1* may contribute to the protective effects of the vaccine on cortical development by attenuating microglial responses in the fetal brains in a maternal immune-activated environment ^106^. Additionally, non-coding *de novo* mutations were demonstrated to affect expression of specific genes, including *Ikzf1*, as a result of altered chromatin interactions ^108^. *Ikzf1* regulates microglial function ^93^, and is expressed in mouse cortical progenitor cells where it plays a key role in differentiation of early cortical neurons ^109^. In summary, these experiments demonstrate the importance of BDNF-TrkB signaling in controlling gene networks associated with regulating social behavior in the LS. The data provides insight about how these signaling pathways may be potentially contributing to etiology and risk for neurodevelopmental disorders that are characterized by changes in social behavior.

## DATA AVAILABILITY

Raw FASTQ sequence data files will be made available from SRA. The code to reproduce the data analyses and figures, along with 1 `SingleCellExperiment` object containing the 70,527 LS nuclei, are available on GitHub https://github.com/LieberInstitute/septum_lateral ^110^. Data is also available in a web-based iSEE app format at https://research.libd.org/septum_lateral/#interactive-websites ^40^. The source data are also publicly available from the Globus endpoint ‘jhpce#septum_lateral’, which is also listed at http://research.libd.org/globus.

## ACKNOWLEDGEMENTS

We thank Linda Orzolek and the Johns Hopkins Single Cell and Transcriptomics Core facility for running sequencing for snRNA-seq libraries. We thank the Joint High Performance Computing Exchange (JHPCE) for providing computing resources for these analyses. We thank Ryan Miller for assisting with data curation. We thank the families of Connie and Stephen Lieber and Milton and Tamar Maltz for their generous support of this work. Schematic illustrations were generated using Biorender and Adobe Illustrator.

## AUTHOR CONTRIBUTIONS

Conceptualization: LAR, SCP, KM

Software: LAR, RGF, HRD, LCT

Formal analysis: LAR, MNT, RGF, AEJ, LCT

Investigation: LAR, EAP, SHK, JHS, YKL, CM, SCP

Data curation: MNT, HRD, LCT

Writing-original draft: LAR, MNT, RGF, SCP, KM

Writing-review and editing: LAR, RGF, LCT, SCP, KM

Visualization: LAR, MNT, RGF

Supervision: LCT, SCP, KM

Project administration: SCP, KM

Funding acquisition: KM, LAR

