## Supplementary figures and images for "TrkB-dependent regulation of molecular signaling across septal cell types"

### Supplementary Figure 1

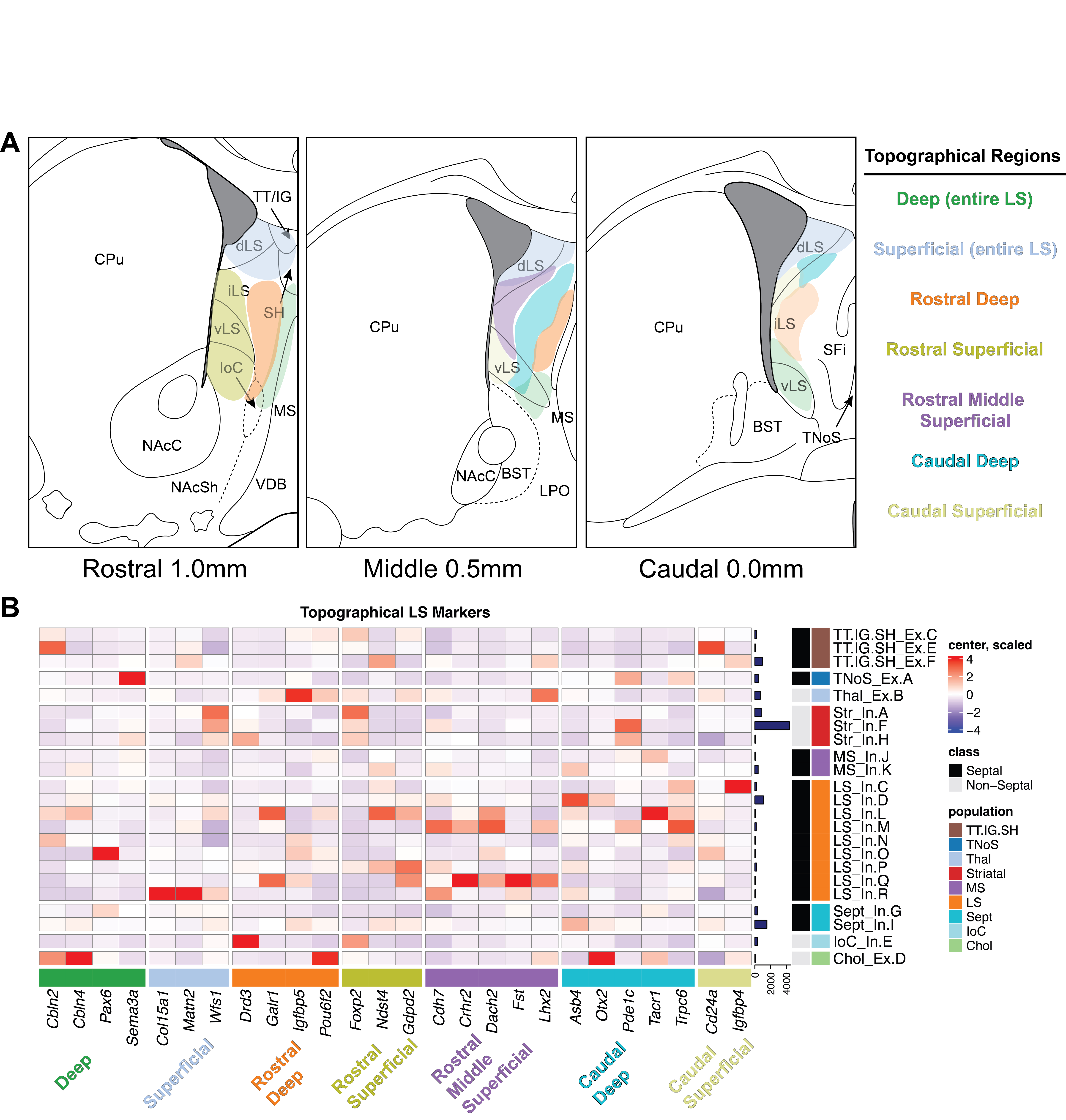

### Supplementary Figure 2

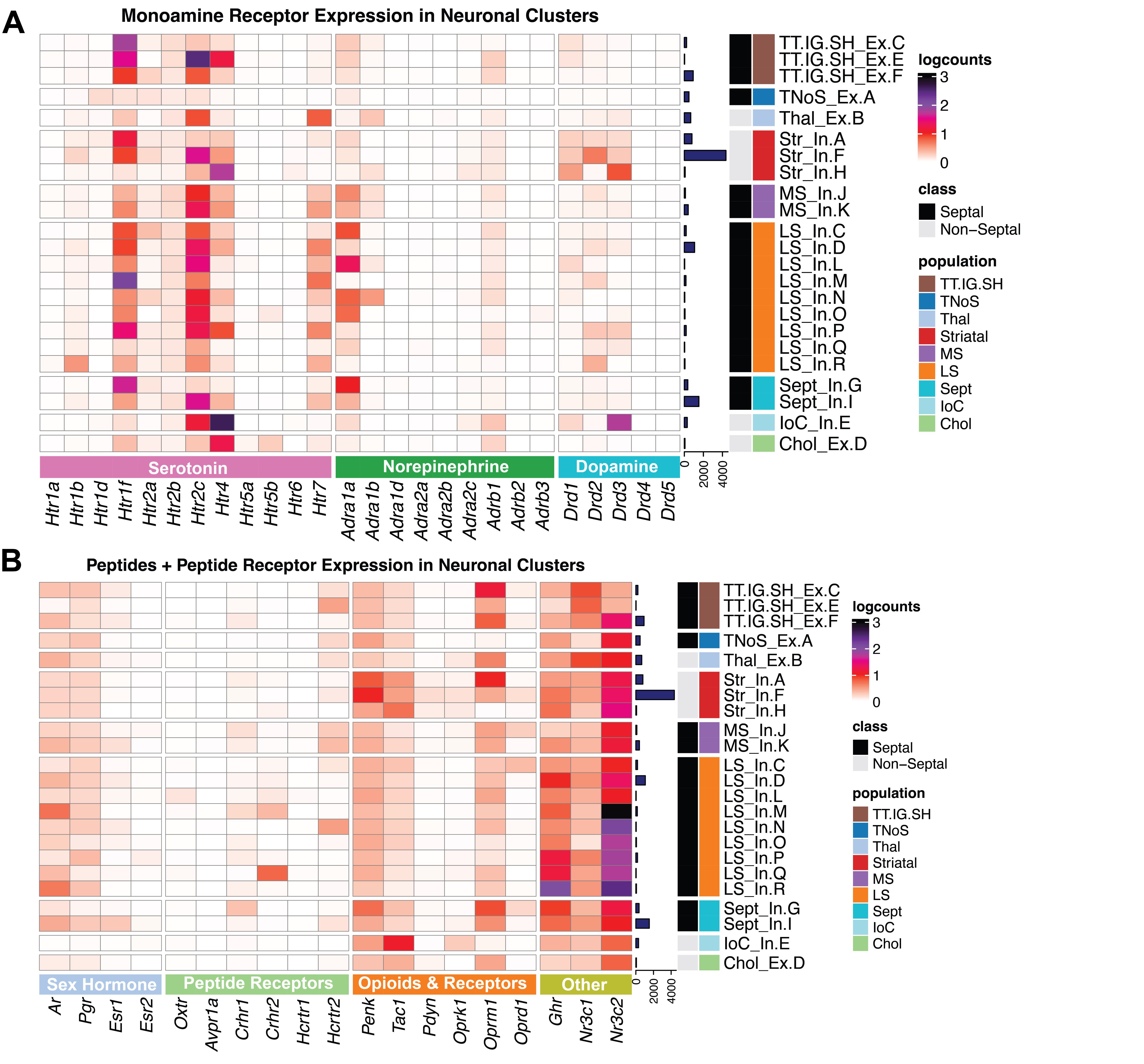
