## Supplemental Text for "TrkB-dependent regulation of molecular signaling across septal cell types"

**Supplementary Figure 1: Septal clusters expressed a wide variety of monoamine, neuropeptides as well as, and neuropeptide and monoamine receptors.** Heatmaps of the (**A**) monoamine receptors and various peptides and (**B**) peptide receptors across neuronal clusters, with normalized expression values (logcounts).

**Supplementary Figure 2: Septal neuronal clusters map onto established LS topographical markers.** (**A**) Illustration depicting the anatomical boundaries across the rostral-caudal axis of the LS, denoting the dorsal (d), intermediate (i), and ventral (v) subregions of the LS, as well as the caudate/putamen (CPu), nucleus accumbens core (NAcC) and shell (NAcSh), islands of calleja (IoC), tenia tecta/indusium griseum (TT/IG), septohippocampal nucleus (SH), medial septum (MS), bed nucleus of the stria terminalis (BST), lateral preoptic area (LPO), triangular nucleus of the septum (TNoS), and septofibrial nucleus (SFi). The illustration is color coded according to the topographical map with distinct genetic markers [^51^](https://sciwheel.com/work/citation?ids=13026226&pre=&suf=&sa=0). (**B**) Heatmap of the genetic markers for the various topographical domains in the LS across the neuronal clusters in the snRNA-seq dataset, with normalized expression values (logcounts) centered and scaled.

**Supplementary Table 1.** A list of the top 100 upregulated DEGs induced by TrkB knockdown in the LS.

**Supplementary Table 2.** A list of the top 100 downregulated DEGs induced by TrkB knockdown in the LS.

**Supplementary Table 3.** A list of the enriched DEGs from the TrkB Knockdown dataset enriched in the microglia cluster and the broad LS cluster.
